# Building on-chip cytoskeletal circuits via branched microtubule networks

**DOI:** 10.1101/2023.10.02.560515

**Authors:** Meisam Zaferani, Ryungeun Song, Sabine Petry, Howard A. Stone

## Abstract

Controllable platforms to engineer robust cytoskeletal scaffolds have the potential to create novel on-chip nanotechnologies. Inspired by axons, we combined the branching microtubule (MT) nucleation pathway with microfabrication to develop “cytoskeletal circuits”. This active matter platform allows control over the adaptive self-organization of uniformly polarized MT arrays via geometric features of microstructures designed within a microfluidic confinement. We build and characterize basic elements, including turns and divisions, as well as complex regulatory elements, such as biased division and MT diodes, to construct various MT architectures on a chip. Our platform could be used in diverse applications, ranging from efficient on-chip molecular transport to mechanical nano-actuators. Further, cytoskeletal circuits can serve as a tool to study how the physical environment contributes to MT architecture in living cells.

**Significance:** Microtubules have essential functions within the cell, including providing a robust railroad for motor-driven cargo transport. The unique properties of microtubules have stimulated attempts to harness these characteristics for targeted delivery of molecular complexes, novel material design, and developing nanotechnologies with precision comparable to living organisms. However, Previous efforts mainly focused on microtubules with fixed length and layout and no controlled MT generation, setting a limit on designing MT architecture. In this study, we integrated nanofabrication with microtubule branching reactions borrowed directly from the cell’s toolkit to construct cytoskeletal circuits and generate microtubule architectures from scratch. That is, our system enables control over microtubule growth and autocatalytic nucleation on a microfluidic chip with micro/nanostructures patterned within.

## Introduction

Microtubules (MTs) are polar biopolymers that grow and shrink (dynamic instability) (1) and perform essential functions within living cells, such as providing tracks along which molecular motors transport cargoes to specific intracellular sites (2, 3). A primary example is the motor-driven transport of molecules and organelles over long distances along axonal MTs (4). Nanotechnology platforms and novel materials have been engineered based on controlling the motility of motors on stabilized MTs (5–8) or organizing static MTs through motor-driven assemblies (9–12). Furthermore, the transport of stabilized MT seeds on surface immobilized motors can be controlled via external forces created by electric (13–15), magnetic (16–18), and optical fields (19–22). These approaches have resulted in development of functional soft active materials (23–27), sensors (28–32), and micro/nanorobots designed for targeted delivery of molecular complexes (15, 33, 34). However, these technologies are constrained by the use of stable MTs with fixed length and thereby limit potential designs and applications.

The next frontier is to harness dynamic MT nucleation and polymerization by directly controlling where MTs are made and in which direction they grow. Such a technology would replicate the cell’s ability to construct a cytoskeletal architecture on demand (35–37), but on a chip rather than within a cell. To achieve such a biomimetic design, it is first necessary to control MT nucleation *in vitro*. Second, MTs must be allowed to grow dynamically. Third, ideally this technology should be composed of MT bundles with uniform polarity growing in a defined orientation to design a robust cytoskeletal architecture. Achieving these three aspects has not been demonstrated thus far. For example, aligning MTs has previously required the use of external forces, which not only requires complicated designs and costly fabrication (38), but also has negative impacts on the structure of MTs, motors, and cargoes. Furthermore, external or depletion forces cannot align MTs into bundles with uniform polarities (39, 40).

It has been discovered that cells have evolved a MT nucleation mechanism to overcome these challenges. In branching MT nucleation, new MTs nucleate at a shallow angle from the side of a pre-existing MT (41), thus conserving the polarity while amplifying the number of MTs. Branching MT nucleation generates a majority of the MTs within the spindle of dividing cells (42, 43). In addition, branching MT nucleation is responsible for generating MTs with uniform polarity in axons (44, 45).

Here, we combined the branching MT nucleation pathway isolated within *Xenopus laevis* meiotic egg extract with microfabrication and microfluidics to build “cytoskeletal circuits”. Our platform mimics the scale and generic shape of axons and enables controlling the active nucleation and growth of branched MT networks for the directed self-organization of MTs within microstructures. Based on this approach we have overcome the three challenges above and engineered tracks of uniformly polarized MTs in a controlled and robust manner. We used quantitative measurements and numerical simulations to elucidate the governing principles of this directed MT self-organizations within various geometric control features. Based on these principles, we finally integrated individual elements to build complex cytoskeletal circuits on a chip.

## Results

To explore how physical boundaries impact branched MT networks, we allowed MTs to grow into sidewalls of a microfluidic reservoir. Branched MT networks were generated in *Xenopus laevis* meiotic egg extract within a reservoir with a width of 200 µm and height of 30 µm (Fig. 1A). Extract was supplemented with Alexa 568-labeled tubulin and GFP-labeled end-binding protein 1 (EB1) to visualize MTs and their growing ends, respectively. Sodium orthovanadate (Vanadate) was used to suppress motor-driven MT gliding on the channel surface, and RanQ69L was used to initiate branching MT nucleation. Upon pipetting 5 µL of the reaction mixture into the reservoir, we used total internal reflection (TIRF) microscopy to observe how branched MT networks formed and interacted with the sidewall of the reservoir.

**Fig. 1.**
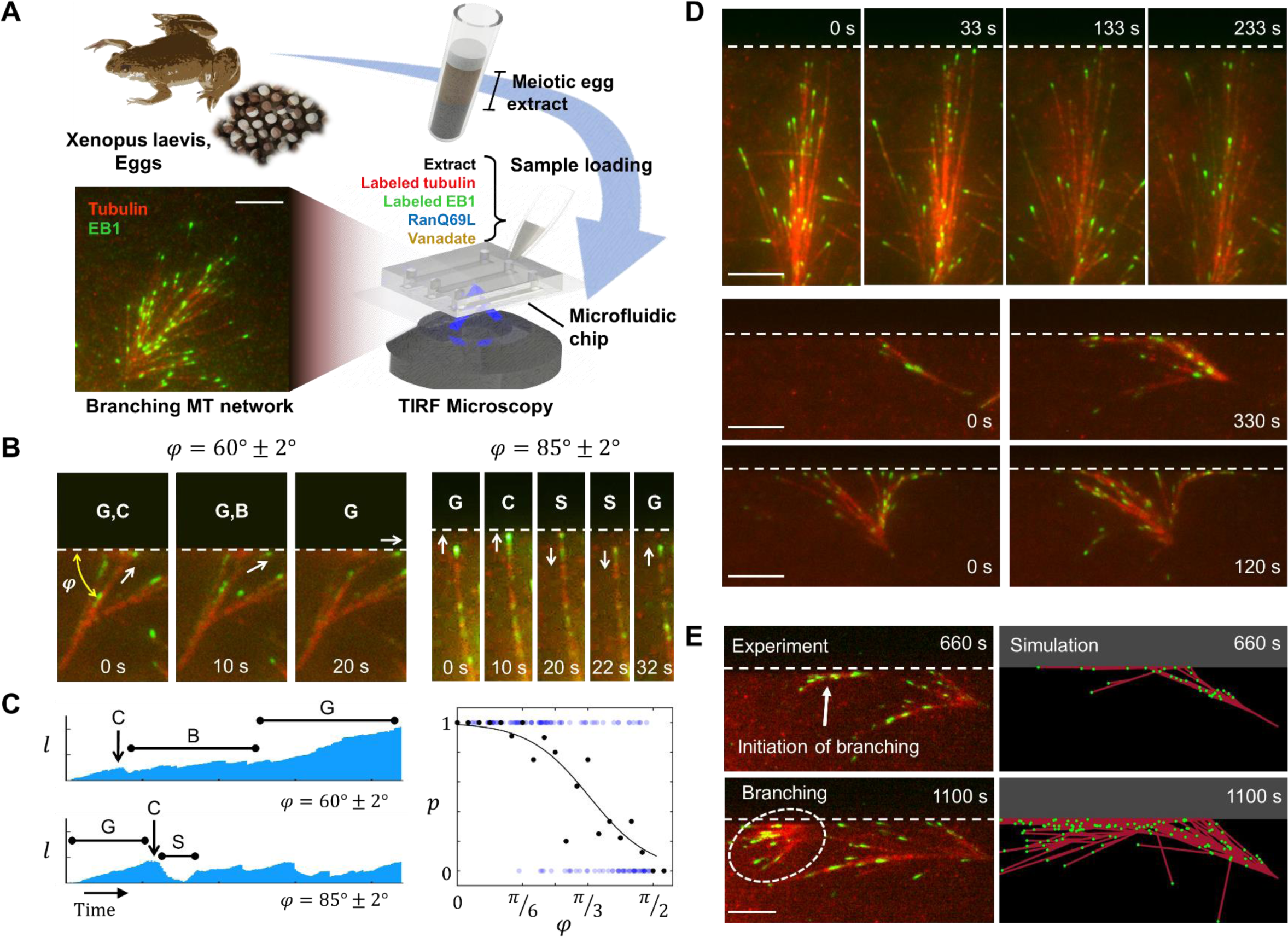
Directed self-organization of branching MTs along a physical boundary. **(A)** Branching MT nucleation within *Xenopus laevis* meiotic egg extract system. The red and green signals in our TIRF imaging system throughout this work represent, respectively, the labeled tubulin to visualize MTs and labeled EB1 proteins to visualize the growing plus end of MTs. **(B)** MT growth dynamics near the sidewall of the microfluidic reservoir depend on their incidence angle with respect to the sidewall (ϕ). G, C, B and S in the images, respectively, denote MT growth, contact (with the sidewall), bending (along the sidewall), and shrinking. **(C) Left:** Kymographs of MT length (*l*) corresponding to MTs shown in (B). **Right:** Probability (*p*) of directed growth along the sidewall vs the angle of incidence. Blue dots correspond to each MT; *p* = 1 when directed growth occurred and *p* = 0 otherwise. Black dots correspond to the probability of directed growth and the solid line is a fitted curve. **(D)** Branched MT networks approaching the sidewall with various distribution of ϕ exhibit different growth dynamics near the sidewall. **(E) Left:** The branching reaction shown in the second row of (D) at a later time. Directed self-organization did not last more than ∼7 min. **Right:** Our stochastic simulations resemble the temporarily directed self-organization of MTs along the sidewall observed in the experiments. Scale bar for all panels: 5 µm.

We observed that as individual MTs from the branched MT networks grow toward and encounter the channel sidewall, their behavior after contact depends on their incident angle (*φ*). For acute angles (Fig. 1B, Movie S1), the growing MTs bend locally (46) at the contact point (Fig. S1A) and continue to grow along the sidewall. In contrast, for angles closer to 90° (Fig. 1B, Movie S2), MTs mostly shrink rapidly (a process known as catastrophe (47)) upon contact with the sidewall. These MTs tended to continue this cycle, i.e., growth, contact, and shrinkage, but we rarely (<10%) observed them to bend and grow along the sidewall throughout the observation period (∼1 hour).

To identify a threshold angle, below which the directed growth most likely occurs, we generated kymographs and plotted the probability of directed growth along the sidewall versus the incident angle (*φ*) (Fig. 1C). By fitting the probability *p*(ϕ) of directed growth to: *p*(ϕ) = (1 + *exp*(α(ϕ − ϕ_*c*_)))^−1^, we identified a threshold for the incidence angle of ϕ_*c*_ = 61° ± 6° (α = 4.1 ± 1.5, with 95% confidence bounds), above which catastrophes are more likely to occur. Not surprisingly, the entire MT network performed directed growth along the sidewall with a similar dependence on the angle with the sidewall (Fig. 1D). A network with MTs mostly growing perpendicular to the wall is not able to grow along the wall (ϕ = 92° ± 6°, Fig. 1D top, and Movie S2), whereas a network approaching the sidewall with acute angles exhibits directed growth along the sidewall (ϕ = 35° ± 10°, Fig. 1D middle, and Movie S1). Consequently, a network with MTs approaching the sidewall with a broad range of angles, including acute and wide angles, can divide on the sidewall and grow in both directions (ϕ = 43° ± 8° and 117° ± 4°, Fig. 1D bottom). Importantly, newly formed MTs that nucleate from the growing MTs along the sidewall also emanate at shallow angles in relation to the sidewall. In other words, the shallow angle between nucleating and existing MTs during the branching process, combined with the directed growth of existing MTs along the sidewall, guides nucleation of new MTs, which we term “directed self-organization” of branched MTs.

The directed self-organization along the sidewall is, however, temporary, as newly forming branched MT networks grow away from the sidewall with shallow angles (ϕ = 17° ± 9°, Fig. 1E left column, Fig. S1B and C and Movie S3). This indicates that, to maintain the directed growth of branched MT networks for long distances and times, one physical boundary is not sufficient, and channels with two sidewalls are required.

To enhance the ability to design directed self-organization of branched MT networks, we developed a computational tool to capture the complex dynamics of these networks as they interact with nearby physical boundaries (Fig. S2). We used our previous computational model and its parameters (e.g., binding rate constant and branching rate constant) as a basis(48) and integrated the probability of directed growth (*p*(ϕ)) obtained in this study. Furthermore, in crowded environments, we hypothesized that MTs require unoccupied space, free from walls or other MTs, for nucleation or growth. Our stochastic simulations consistently reproduced the experimentally observed behavior of MTs with sidewalls (Fig. 1E right column, Movie S3). Thus, this computational tool can be used to predict the self-organization of branched MT networks in more complex geometric configurations, as pursued in conjunction with experiments next.

To test whether we could maintain the directed self-organization of branched MT networks by two sidewalls within a narrow channel (Fig. 2A), we fabricated photoresist-based microstructures on a coverslip glass with a main channel, which is in contact with the rest of the microfluidic reservoir only from the top and one opening from the side (Fig. 2A and Fig. S3B). The extract reaction mixture was pipetted into the reservoir to initiate the reaction and imaged via TIRF microscopy. Branched MT networks first organized in the reservoir, before entering the side openings of the channels, where they propagated (Fig. 2B and Fig. S3C). We visualized their self-organization within straight channels with a width of 2 µm and lengths of 15 µm, 30 µm, and 60 µm (Fig. 2C). We observed a persistent directed self-organization of MTs inside these channels with a growth rate of 6.9 ± 2.0 μm/min (Fig. S4A-D) such that the growth over 60 µm length occurred within approximately 10-15 minutes after MT entry (∼ 45-60 min from extract injection) to the channels (Fig. 2C bottom). We also examined how the width of the channel impacts the self-organization of MTs (Fig. 2D and Movie S4). By tracking EB1 spots and extracting MT growth trajectories within the channels, we found that MTs grow in straight lines (Fig. S4E-F) and the distribution of growth directions becomes narrower in channels with smaller widths, from 2 µm to 0.5 µm (Fig. S4A and B). Further, we noticed that MTs aligned with the channel grew for longer distances before they shrink (Fig. S4C). By counting the number of EB1 spots in the channels with lengths of 30 µm and widths of 0.5, 1, and 2 µm, we found that the number of growing MTs in the channels increased over time, reaching a fluctuating plateau within approximately 10 minutes from their entry to the channel (Fig. 2E). This observation, which was similarly observed in our simulations, suggests that the maximum number of growing MTs in channels with fixed lengths, i.e., channel capacity, depends on the width of the channel (Fig. 2F and Fig. S4G-K). Note that besides changing the channel geometry, the number of MTs can be modulated by RanGTP. As expected, increasing the concentration of RanGTP also increases the number of MTs in the channel (Fig. 2G).

**Fig. 2.**
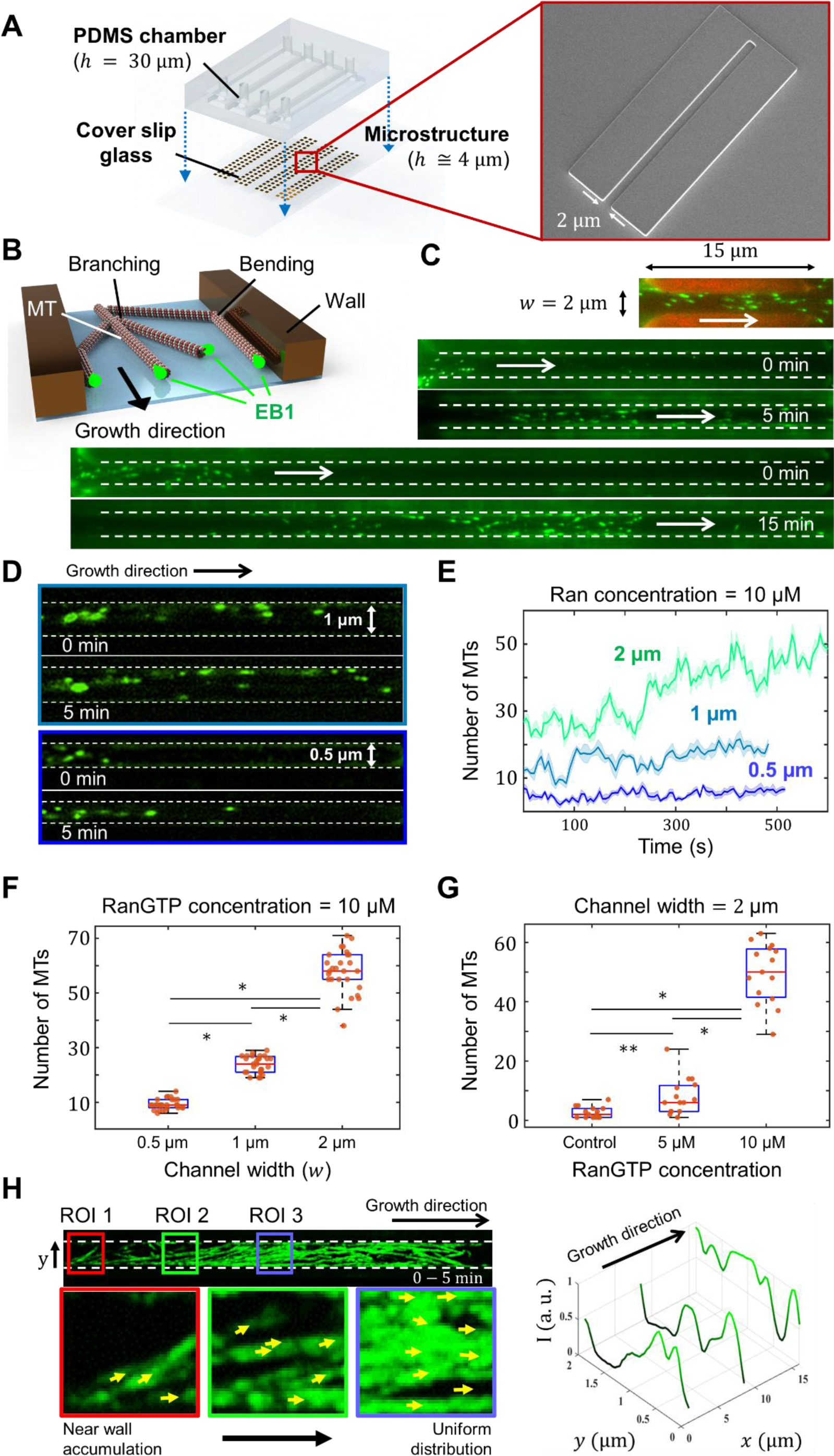
Self-organization of branching MTs within straight channels. **(A)** Schematic (left) and scanning electron microscopy image (right) of the microstructures fabricated on the bottom surface of the channel. **(B)** Schematics and **(C)** experimental observations of branched MT networks self-organizing within straight channels with 15, 30 and 60 µm lengths. White arrows show the growth direction of MTs **(D)** Branched MTs within channels with 30 µm length and 1 µm (top) and 0.5 µm (bottom) widths. **(E)** The number of EB1 spots (growing MTs) increases over time and reaches a plateau determined by the channel width. **(F)** Number of MTs in the channel can be controlled by the channel width and **(G)** RanGTP concentration. * p-value < 0.00001, ** p-value < 0.001. **(H)** The EB1 signals overlapped (1 fps) for 5 min and the corresponding normalized light intensity (I) across the channel width at different ROIs. More uniform distribution of intensity *I* across the channel width indicates a more uniform distribution of EB1 spots.

It is important to note that the spatial distribution of MTs across the channel width also changes as the MT network propagates within the channel. We quantified this observation by measuring the intensity profile of EB1 spots (overlapped with 1 fps over 5 min) along the channel width and found that branched MTs become more uniformly distributed across the channel width as they grow inside the channel (Fig. 2H and Fig. S5).

To further investigate the relationship between channel capacity and width, we examined the self-organization of MTs within a channel featuring varying widths (Fig. 3A and Movie S5). In this experiment, the channel comprised three successive compartments with widths of 2 µm, 1 µm, and 2 µm, respectively. By counting the number of growing MTs (i.e., via EB1 spots) within certain regions of interest (ROI) (2 µm × 4 µm, Fig. 3A), we observed a decrease in the number of MTs as they self-organized within the narrow region (Fig. 3B, C). Nevertheless, we did not observe an increase in the number of growing MTs as they entered the third compartment, which had a length of 5 µm (Fig. 3C). In contrast, when we increased the length of the third compartment to 15 µm (Fig. 3D and Movie S5), we observed branching MT nucleation occurring within the third compartment (Fig. 3E top). Consequently, the newly generated MTs adapted the mass of the network to the channel capacity in the third compartment (Fig. 3E bottom). This length-sensitive onset of branching nucleation is most likely due to the time and length required for MTs to recruit the branching factors. This observation, which was supported by our simulation predictions (Fig. S6), indicates that, subject to sufficient space and time, autocatalytic branching nucleation(49) leads to dynamic adaptation of MT numbers in response to the geometric characteristics of the channel.

**Fig. 3.**
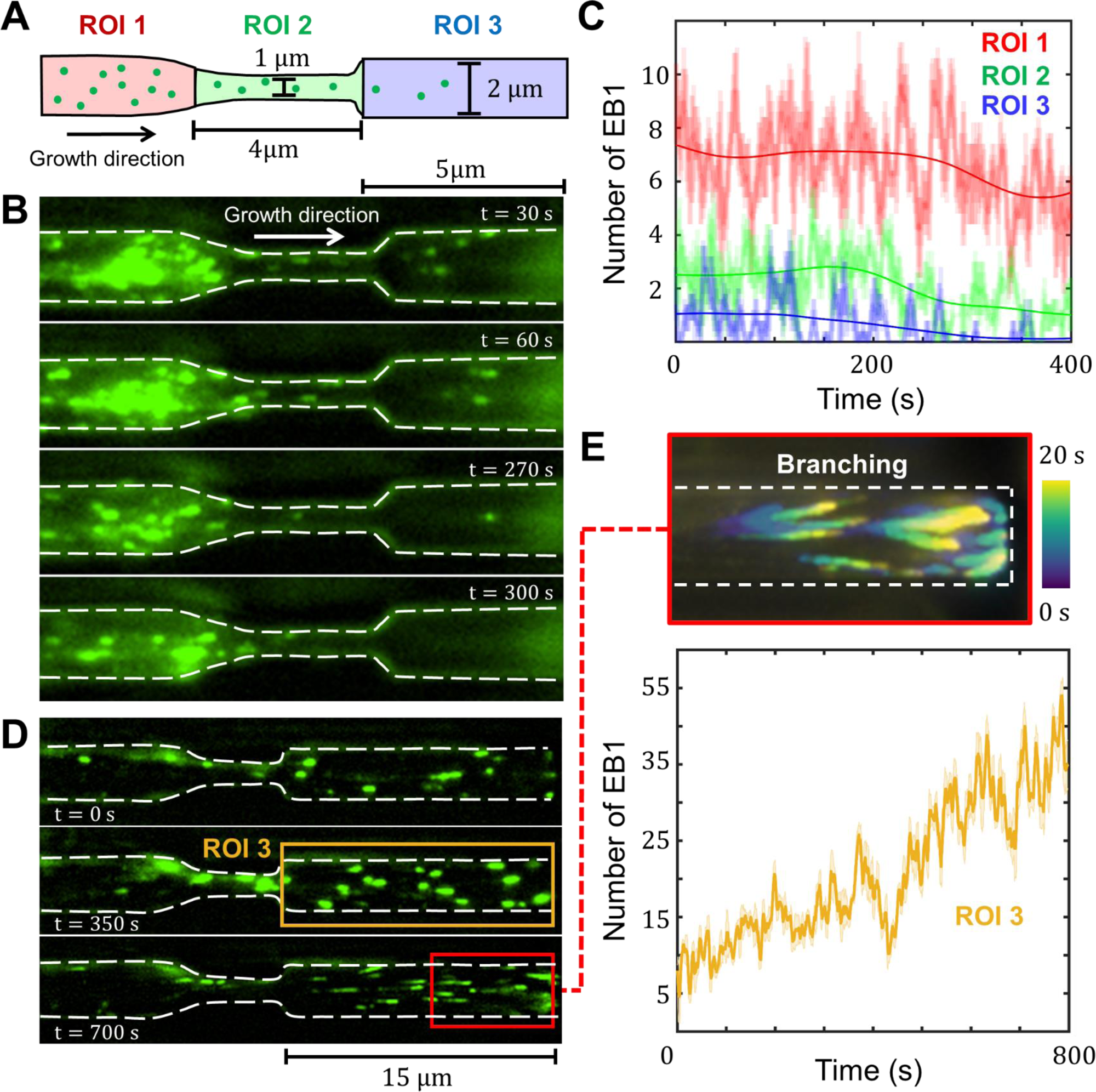
Adaptive self-organization of MTs through channels with varying width. **(A)** Schematic of a channel composed of three compartments with different widths. **(B)** Branching and growing MTs self-organize and adapt their number as they grow through the narrow channel. **(C)** Number of EB1 spots over time. The number of EB1 in the third compartment does not correspond to its width. **(D)** A channel similar to (B), but the length of the third compartment has been increased to 15 µm. **(E)** The EB1 signals overlapped (5 fps) for 20 s and color coded according to the elapsed time (top). Branching nucleation occurs and the number of EB1 increases over time to adapt the growing MTs number with the channel width (bottom).

Based on how MT networks behave in straight channels, the next pressing question was how branched MT networks behave in more complex patterns. We constructed channels with symmetric and asymmetric bifurcations and observed how branched MT networks behave upon encountering them (Fig. 4A, Figs. S7A, S8A and B, and Movie S6). By probing the growing MT flux defined as the average number of EB1 spots per seconds within a certain ROI (Materials and Methods), we found that symmetric bifurcations resulted in, on average, an equal growing MT flux in each division (ℐ_1_ = 7.2 ± 1.3 and ℐ_2_ = 6.8 ± 1.4). In contrast, asymmetric bifurcations produced fewer MTs growing within the channel branching to the side (ℐ_1_ = 8.7 ± 1.1 and ℐ_2_ = 4.3 ± 0.3, Fig. 4A and Fig. S8A and B). To further understand the principles of MT network division, we quantified growth in a hepta-furcated geometry. In this configuration, which resembles the generic shape of a neuronal growth cone, equal MT fluxes on average were observed inside channels designed symmetrically with respect to the main channel (Fig. 4B, Fig. S8C and D, and Movie S6). However, more MTs grow inside the side channels with smaller angles.

**Fig. 4.**
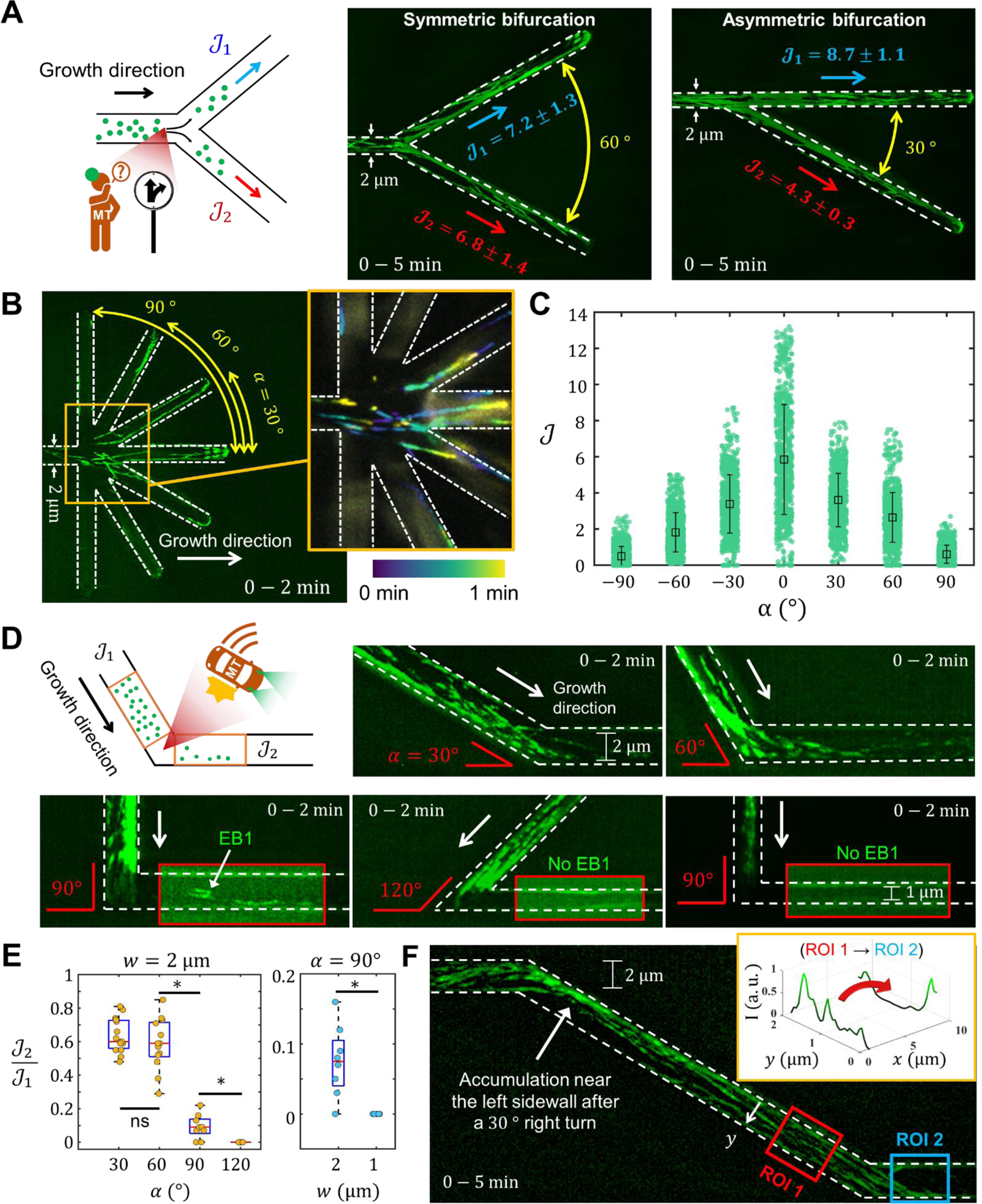
Self-organization of branched MT networks through geometrical divisions and turns. **(A)** Division of branching MT networks through symmetric and asymmetric bifurcations (channel width is 2 µm). The average number of EB1 spots per second (ℐ), were measured to quantify the division process. The experimental images are overlapped EB1 signals acquired for 5 min at 1 fps rate. **(B)** Experimental quantification of MT network divisions through a hepta-furcated geometry. Image acquisition setup is similar to (A). **(C)** The average number of EB1 spots per second in each side channel of the hepta-furcated geometry. **(D)** Bending of branching MT networks through turns with different angles in the channel (channel width is 2 µm). Sharper turns result in fewer MTs able to bend and continue to self-organize through the turn; a 90° turn results in loss of more than 90% of MTs during the turn. This number further decreased to 0 when the channel width was 1 m (bottom right). **(E)** We measured the average number of EB1 spots per second before and after the turn, which yields the fractional loss of MTs. Loss of MTs depends on both turning angle (left) and channel size (right). * p-value <0.001. **(F)** Turns in the channel shift the spatial distribution of EB1s. Overlapped EB1 signals acquired for 5 min at 1 fps from MTs in a channel featuring two consecutive right and left turns. As quantified by the light intensity profile across the channel width, each right/left turn results in a corresponding shift of EB1 spots toward the left/right sidewall.

To guide MT networks in more complex patterns, it is necessary to propagate MTs through turns in the channels. We therefore investigated the self-organization of branching MTs in curved channels by analyzing MT growth in 2 µm-wide channels featuring 30°, 60°, 90°, and 120° turns (Fig. 4D, Fig. S8E, and Movie S7). By measuring MT flux before and after each turn, we quantified the loss in the number of EB1 spots associated with each turn. This loss depends on the turning angle (Fig. 4E left and Fig. S7B), i.e., MTs growing more perpendicular to the sidewall encountered at the turn were more likely to catastrophe and thus less likely to bend and grow along the sidewall. This is consistent with results shown in Fig. 1C. Interestingly we observed approximately 10% of MTs bending and growing through a 90° turn. However, this number decreased to 0 when the channel width was reduced to 1 µm, where the distribution of MT orientations approaching the curved sidewall was narrower and closer to 90° (Fig. 4D bottom right, Fig. 4E right). To achieve a thorough characterization, we conducted simulations to quantify how the combination of turning angle and channel width determines the loss of MTs (Fig. S8F). Although each turn resulted in a loss of MTs, we found that branching MT nucleation increases MT number after each turn and adjusts it to the channel capacity within 7 ± 1 µm for 30°, 9 ± 2 µm for 60°, and 14 ± 4 µm for 90° turns (Fig. S9). Therefore, branching MT nucleation not only dynamically adapts the number of MTs according to the channel width but also compensates for the loss of MTs, thereby promoting the formation of a robust MT architecture.

More importantly, we observed that growth of MTs on straight lines results in a temporary accumulation of growing MTs along one of the sidewalls after each turn in the channel: right turns resulted in MT accumulation near the left sidewall and vice versa (Fig. 4F). Therefore, incorporating turns in a channel is a useful technique for temporarily modulating the spatial distribution of MTs across the channel width and building regulatory elements in the circuit.

A fundamental regulatory function involves controlling the division of branched MT networks within splitting channels. Although we found that geometrical characteristics of the splitting channels govern the division of branched MT networks, modulating the spatial distribution of MTs across the channel width before dividing them can control the MT division process. Based on our simulations (Fig. 5A and Movie S8), we proposed that a turn in the channel before a symmetric bifurcation can lead to an asymmetric division of branched MT networks. For instance, a left turn before symmetric division can effectively guide MT growth toward the right arm of the bifurcation, thus regulating MT divisions.

**Fig. 5.**
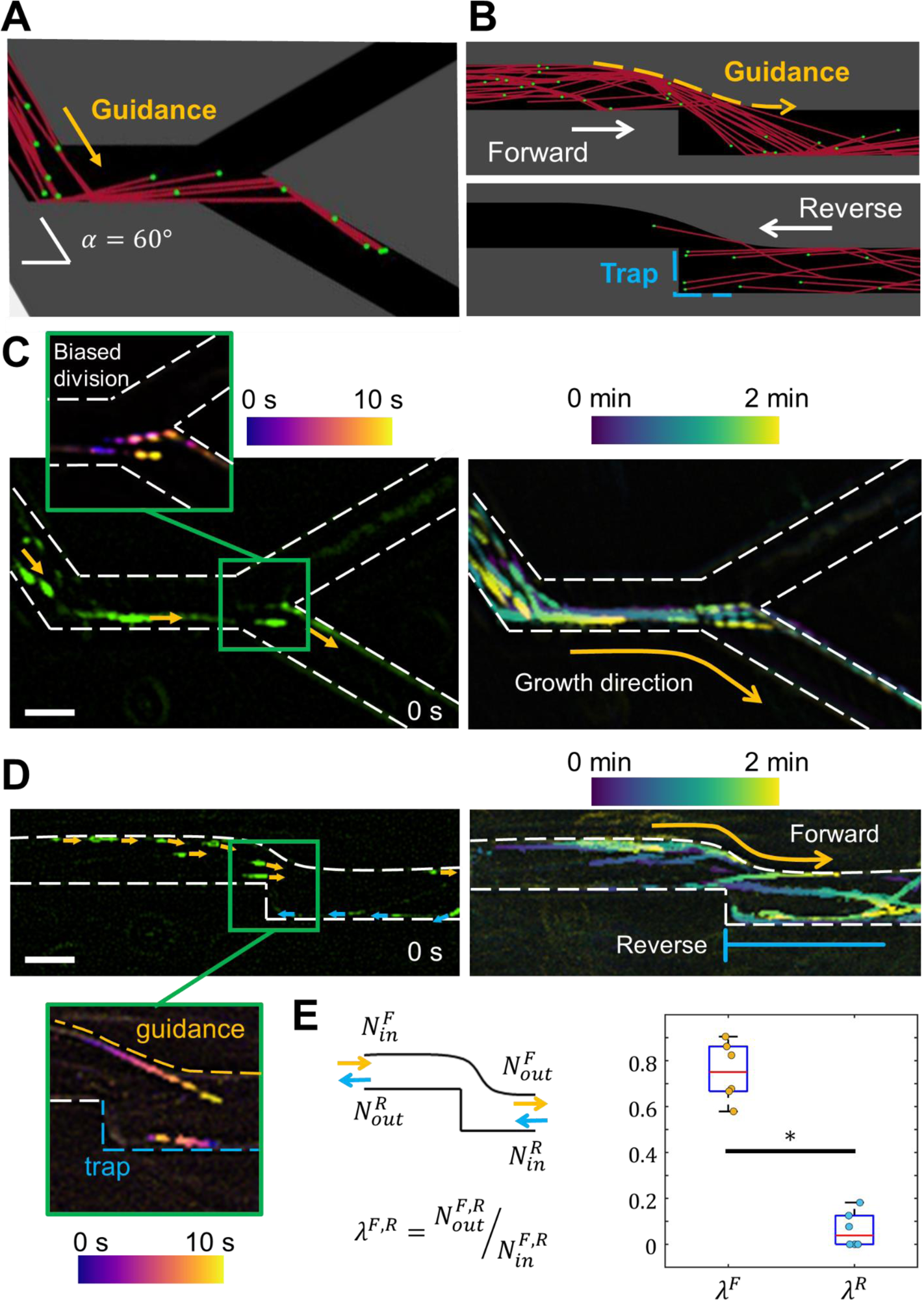
Modulation of MT self-organization to build regulatory elements. **(A)** Our simulations of a symmetric bifurcation preceded by a turn led to an asymmetric division of MT networks. **(B)** Simulation of a MT diode. Self-organization of MTs is allowed in the forward direction and hindered in the reverse direction. **(C)** Experimental realization of biased division. For better visualization, EB1 signals are overlapped at 1 fps and color coded according to the elapsed time. Scale bar: 2 µm. **(D)** Experimental realization of a MT diode. Imaging setup is similar to (C). Scale bar: 2 µm. (E) Function of MT diode was quantified for 6 replicates to confirm the existing bias in the MT growth direction. * p-value < 0.00001.

Another useful regulatory element is a “MT diode”, which can select for the polarity of MTs within a channel. Such a channel would permit MTs whose polarities align with the forward direction of the channel to grow and reach the other end of the channel, while hindering the growth of MTs with polarities aligned in the reverse direction. Our simulations indicated that a channel with a smoothly-turning left sidewall and a 90°-turning right sidewall can induce directional bias in MT growth, resulting in a functioning MT diode. In this configuration, MTs accumulated along the left sidewall and growth in the forward direction can continue, while those accumulated along the right sidewall and growing in the reverse direction become trapped behind the turn (Fig. 5B and Movie S9). Note that the accumulation of MTs along the left and right sidewalls is induced, respectively, via preceding 30° turns to the right and left within the channel.

To confirm these predictions, we experimentally probed the self-organization of MT networks within the proposed microstructures and observed both biased-division (Fig. 5C and Movie S8) and selection for the polarity within the MT diode configuration (Fig. 5D and Movie S9). To quantitatively assess the function of our MT diode (Fig. 5E), we measured the fraction of MTs self-organized through the channel in the forward direction (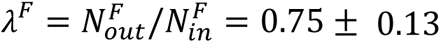) and found it significantly greater than the same fraction in the reverse direction (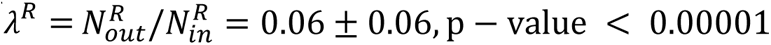). As such, our proposed regulatory elements function properly and regulate the self-organization of MTs as predicted by simulations.

## Discussion

In summary, we have developed cytoskeletal circuits that serve as a platform for engineering robust MT architectures with uniformly polarized MTs generated via the branching nucleation pathway. In this active matter platform, the self-organization of branching MTs is controlled by the geometrical features of microstructures fabricated inside a microfluidic chamber. We established the basic principles of this self-organization and utilized these principles to construct regulatory elements as examples, showcasing the potential of cytoskeletal circuits in designing complex MT architectures.

Our cytoskeletal circuits have the potential to create numerous on-chip nanotechnologies for exploration. We propose employing biomolecular motors, such as kinesin-1, within the cytoskeletal circuits to develop robust on-chip molecular transport on branching MTs. The intrinsic uniform polarity and high density of MTs within cytoskeletal circuits potentially will allow for high-yield molecular transport with efficiencies comparable to those of living systems. Additionally, cytoskeletal circuits could be expanded by integrating them with other chemical, optical, magnetic, and electrical components to create more complex and dynamic structures, such as cytoskeletal switches and logic gates.

Although our primary focus was on MT architecture within cytoskeletal circuits, it is worth noting that the tips of MTs can generate forces as they grow(50–52). Consequently, the directed force generated by branching MTs in cytoskeletal circuits could serve as the foundation for developing miniature actuators and manipulating micron/nano-sized objects. Furthermore, the unique mechanical properties of MTs can allow for nano-scale mechano-sensing within cytoskeletal circuits(53–55). In a more fundamental context, our cytoskeletal circuits could also be utilized to investigate how physical environments within the axon, as well as MT-associated proteins such as TPX2, can contribute to axonal MT architecture(45).

## Materials and Methods

### (I) Protein Purification and Xenopus egg extract

#### Purification of EB1-GFP

To purify EB1-GFP, the following established methods were employed(56). Initially, the protein was expressed in *E. coli* (Rosetta 2 strain) at 37°C for 4 hours. Subsequently, cellular lysis was accomplished using an EmulsiFlex (Avestin) in lysis buffer composed of 50 mM NaPO4 (pH 7.4), 500 mM NaCl, 20 mM imidazole, 2.5 mM PMSF, 6 mM CME, 1 cOmpleteTM EDTA-free Protease Inhibitor (Sigma), and 1000 U DNAse 1 (Sigma). Affinity chromatography utilizing a HisTrap HP 5 mL column (GE Healthcare) was then employed to purify the protein in binding buffer consisting of 50 mM NaPO4 (pH 7.4), 500 mM NaCl, 20 mM imidazole, 2.5 mM PMSF, and 6 mM BME. Elution of the protein was carried out using an elution buffer containing 50 mM NaPO4 (pH 7.4), 500 mM NaCl, 500 mM imidazole, 2.5 mM PMSF, and 6 mM BME. The resulting peak fractions were pooled and loaded onto a Superdex 200 pg 16/600 gel filtration column. Gel filtration was performed in CSF-XB (10 mM HEPES, pH 7.7, 1 mM MgCl2, 100 mM KCl, 5 mM EGTA) supplemented with 10% (w/v) sucrose.

#### Purification of RanQ69L

RanQ69L, utilized to induce branching MT nucleation, was purified following previously described procedures(41, 57). The RanQ69L variant, with an N-terminal BFP tag for enhanced solubility, was expressed and lysed using the lysis buffer (100 mM tris-HCl, pH 8.0, 450 mM NaCl, 1 mM MgCl2, 1 mM EDTA, 0.5 mM PMSF, 6 mM BME, 200 µM GTP, 1 cOmpleteTM EDTA-free Protease Inhibitor, 1000 U DNAse 1). Affinity purification of the protein was achieved using a StrepTrap HP 5 mL column (GE Healthcare) in binding buffer consisting of 100 mM tris-HCl, pH 8.0, 450 mM NaCl, 1 mM MgCl2, 1 mM EDTA, 0.5 mM PMSF, 6 mM BME, and 200 µM GTP. Elution of the bound protein was performed using elution buffer composed of 100 mM tris-HCl, pH 8.0, 450 mM NaCl, 1 mM MgCl2, 1 mM EDTA, 0.5 mM PMSF, 6 mM BME, 200 µM GTP, and 2.5 mM D-desthiobiotin. Finally, the eluted protein was dialyzed overnight into CSF-XB (10 mM HEPES, pH 7.7, 1 mM MgCl2, 100 mM KCl, 5 mM EGTA) containing 10% (w/v) sucrose.

Tubulin from bovine brain (PurSolutions) was also labeled with Alexa 568 NHS ester (GE Healthcare) as previously described(58).

#### Xenopus egg extract

The branching MT nucleation reactions in the Xenopus egg extract system followed established protocols(41, 59). Briefly, Xenopus cytosol, naturally paused in meiosis II(60, 61), was supplemented with fluorescently labeled tubulin (at a final concentration of 1 µM) to visualize MTs. GFP-labeled EB1 (at a final concentration of 100 nM) was also added to track MT plus ends. Additionally, RanQ69L (at a final concentration of 10 µM) was included to induce branching nucleation, while sodium orthovanadate (at a final concentration of 0.5 µM) was incorporated to inhibit motors and prevent MT gliding on the bottom surface of our microfluidic chamber. The reaction mixture was prepared and kept on ice until it was injected into the microfluidic chamber.

### (II) Microfabrication process

Microfluidic chips were fabricated by bonding a polydimethylsiloxane (PDMS) block (Sylgard 184, Dow Corning) with microchannels and microstructure-featured coverslip glass, as illustrated in Fig. 2A and Fig. S3. First, the coverslip glasses (No. 1.5, CG15KH1, Thorlab) were cleaned using sonication in 70% ethanol for 20 min., followed by rinsing with DI water and drying with nitrogen gas. Subsequently, the glasses were completely dried in a convection oven at 70 °C. To enhance chemical adhesion of the photoresist to the coverslip glasses, the glasses were dehydrated and primed with hexamethyldisilane (HMDS) at 148°C in a YES vacuum oven (Yield Engineering Systems). Next, microstructures with a thickness of ∼4 μm were patterned onto the HMDS-primed surface using AZ 4330 photoresist (MicroChemicals). The photoresist was spin-coated at 4000 rpm for 40 s and soft-baked at 110°C for 60 s. A high-resolution pattern generator (DWL 66+, Heidelberg instruments) was utilized to expose the photoresist to 405 nm UV light. After exposure, the photoresist structures were developed for 3 min. in the AZ300MIF developer (MicroChemicals), and the remaining developer was rinsed using DI water and dried with nitrogen gas.

The fabrication of the microchannel containing PDMS blocks was carried out using standard soft-lithography techniques. Initially, a 4-inch silicon wafer was spin-coated with SU-8 2025 photoresist (MicroChemicals) at 3000 rpm. The coated wafer underwent a soft-baking process on a hot plate at 65℃ for 1 min and 95℃ for 5 min. Next, the SU-8 photoresist was exposed to 375 nm UV light using a medium-resolution pattern generator (μPG 101, Heidelberg instruments) to generate channel patterns. Then it was post-exposure-baked on a hot plate at 65°C for 1 min and 95°C for 5 min. before being developed in a SU-8 developer (MicroChemicals) for 5 min. After development, the wafer was rinsed with isopropyl alcohol, dried with nitrogen gas, and then hard-baked on a hot plate at 120℃ for 30 min. PDMS mixture consisting of a 10:1 weight ratio of PDMS base to curing agent was poured on the SU-8 mold, degassed to remove bubbles, and baked in a convection oven at 70℃ for 5 hours. The cured PDMS was then peeled off from the mold, cut into blocks containing microchannels, and punctured to make inlet and outlet.

To bond the coverslip glass and PDMS, both were functionalized under air plasma treatment via plasma cleaner (PDC-002, Harrick Plasma). The treated surfaces were carefully aligned and then pressed together. The bonded chips were baked in a convection oven at 70℃ for 30 min. to enhance the bonding strength between the PDMS and coverslip glass. Each microfluidic chip consisted of four 1.6 × 10 mm microchannels, and an approximately 30 × 50 microstructure array was placed within each microfluidic chamber.

### (III) Microscopy and Image processing

A Nikon TiE microscope equipped with a 100 × 1.49 NA objective was used for Total Internal Reflection Fluorescence (TIRF) microscopy. Image capture was done using an Andor Zyla sCMOS camera. The NIS-Elements software (Nikon) was utilized for the acquisition process to capture dual-color images. For optimal visualization, adjustments were made to the brightness, contrast, and acquisition rates (0.2, 1, and 5 fps) for each experiment.

To measure the kymographs and curvature of microtubules (MTs) as shown in Fig. 1 and Fig. S1, we manually extracted their shape at each frame by assigning approximately 100 points (Xi, Yi) to their image in the tubulin channel. After coarse graining the trajectories, we calculated the MT curvature and arc length at each frame using the equations:

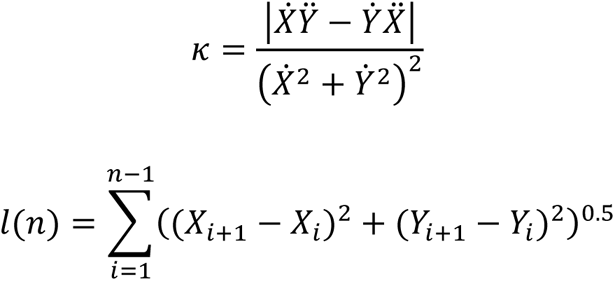

Note that dots denote time-derivatives.

To detect EB1 comets on the MT plus ends, we first removed the background from the EB1 channel signal and applied a Gaussian filter with a standard deviation of 2 pixels. We then used TrackMate to detect EB1 spots and obtain trajectories of MT growth using a nearest neighbor algorithm(62). The parameters in TrackMate were optimized for each dataset to minimize errors in the analysis. Additionally, we manually verified the tracks to ensure their accuracy. These trajectories were used to calculate the MT’s growth rate, mean-square displacement (MSD), velocity autocorrelation correlation, processive length, and growing angle using custom MATLAB codes (Fig. 2, S4, and S6):

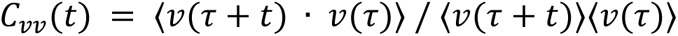

To measure the MT flux in each region of interest (Fig. 3, 4, and S8), we counted the number of EB1 comets in each frame and averaged them over a 10-second time window to smooth out fluctuations associated with EB1 detection:

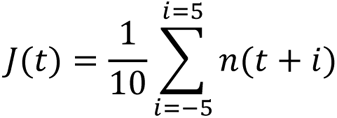

### (IV) Numerical simulation

#### Initial conditions and parameters of simulation

Stochastic kinetic simulations of branched MT networks and their interaction with structures were computed by MATLAB (R2021b, MathWorks). The scenario of MT branching and related parameters, such as branching angle distribution, binding rate constant for deposition of nucleation sites (*k*_*bind*_ = 0.1 molecules/μm · s) and branching rate constant (*k*_*branch*_ = 2.5 × 10^−4^ s^−1^/molecule), were obtained from the sequential reaction model of previous research(48). The remaining parameters, such as growth speed of MT plus-ends (*v*_*pe*_ = 6.9 μm/min) and probability of directed growth along the sidewall (*p*(ϕ) = (1 + exp(α(ϕ −ϕ_*c*_)))^−1^), where α = 4.1 and ϕ_*c*_ = 61°), were derived from the experimental results shown in Fig. 1C and Fig. S4D. The simulation was initiated at *t* = 0 s with MTs of 25 nm in diameter. The minus-ends of each MT were randomly distributed within an initial region, and all subsequent MTs were nucleated solely through branching from existing ones.

#### Stochastic model for self-organization of branching MT networks

A microtubule is modeled as a series of rectangles with 25 nm width. For every time step (Δ*t* = 0.5 s), the plus-end position (*s*_*pe*_) and growth direction (θ) of all MTs were computed iteratively according to their growth state: (i) nucleation/branching, (ii) growth, (iii) catastrophe. Detailed scenarios for each growth state are as follows:

##### (i) nucleation/branching

The scenario for nucleation of new branched MTs benchmarked the sequential reaction model as illustrated in Fig. S2A. The new branching molecule 1 binds on the existing MT when *t*_*bind*_ = *t*_0_ + exprnd(1/*k*_*bind*_)/∫ *ds*_*pe*_, where exprnd is the MATLAB function to generate exponential random numbers and *t*_0_ is the nucleation time of the mother MT. The binding site of branching molecule 1 is defined randomly along the mother MT length. Branched MTs are then nucleated from these bound, inactive sites after *t*_*branch*_ = exprnd(1/*k*_branch_) due to branching molecule 2. In the sequential reaction model, the spatial occupation by individual MTs was not considered, allowing any branching complex to nucleate a new MT. However, in this model, it was assumed that branching complexes can only bind and nucleate a new MT if the binding sites are unoccupied by walls or other MTs, as illustrated in Fig. S2B and C. Therefore, the presence of other MTs or walls in the binding space of the branching complex, assumed to have a radius of 50 nm(63), is checked. A MT is only nucleated when the surrounding area is unoccupied. The branch angle was selected from a Gaussian distribution with a mean of 0° and a standard deviation of 9° (48), and then the growth state of the new MT is shifted to (ii) growth.

##### (ii) growth

MTs are considered to have a fixed minus-end and grow at a prescribed growth speed (*v*_*pe*_) at the plus-end. Therefore, the position of plus-end (*s*_*pe*_) is updated at each time step as

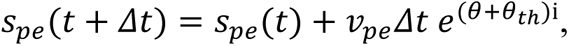

where θ_*t*ℎ_is the temporal tilting angle to simulate the thermal fluctuation of MT, and **i** denotes an imaginary unit. In the real system, MTs fluctuate in various bending modes(64), but we only consider the first bending mode to simplify the simulation model. Then the tilting angle θ_*t*ℎ_(*S*) over the arc length of MT (*S* = ∫ *ds*_*pe*_) constrained to a plane can be described as ⟨cos θ_*t*ℎ_(*S*)⟩ = exp(−*S*/*L*_*p*_), where *L*_*p*_is the persistence length of MTs and the angle brackets denote an average value(65). Keller *et al.* reported the statistics of the deflection ratio and persistence length of MTs in *Xenopus laevis* egg extract(66), thus their data was utilized to estimate the thermal fluctuation-driven random trajectories of MTs.

When a growing MT encounters a microstructure wall, it undergoes bending parallel to the wall, taking on the direction that results in the minimum incidence angle (ϕ). To determine whether the MT continues to grow or undergoes shrinking, the probability of directed growth, *p*(ϕ), is compared to a random number (0 ≤ ψ ≤ 1). If *p*(ϕ) ≥ ψ, the MT bends and continues to grow. However, if *p*(ϕ) < ψ, the MT cannot grow, and its growth state transits to (iii) catastrophe (Fig. S2B).

In this model, individual MTs occupy their own non-overlapping space, limiting the number of MTs that can be positioned inside the channel as observed in Fig. 2. The interaction between MTs in this crowded environment is still unclear, but assuming a scenario depicted in Fig. S2C yields reasonable results, as confirmed in Fig. S5. When a growing MT encounters another MT, its behavior varies based on the incidence angle. At low incidence angles, the MT bends and forms a bundle with the encountered MT. At high incidence angles, the growing MT crosses over the encountered MT(54, 67). In crowded environments, a growing MT can encounter a MT bundle, and in such cases, even if it crosses over one MT, it may continue to encounter additional MTs within the bundle. Therefore, when encountering an MT bundle, it is assumed that the growing MT bends and forms a bundle regardless of the incidence angle. This scenario was implemented using a simple approach. When a MT encounter other MTs, it grows towards the point within a certain radius where the bending angle (ϕ_*bend*_) is minimized. The radius was set to 150 nm to match the simulation and experimental results. Similarly, to encountering a wall, if *p*(ϕ_bend_) is smaller than ψ, the MT cannot continue growing and its growth state transitions to (iii) catastrophe.

##### (iii) catastrophe

In this state, the length of an MT shrinks to zero and inactive branching molecules bound on this MT are removed. It is assumed that already nucleated MTs continue to grow even if the mother MT undergoes catastrophe.

## Data and code availability

The code used to compute the simulations of this work is available at https://doi.org/10.5281/zenodo.8105932. All other data are available in the main text or the supplementary materials.”

The main simulation code is provided in the file *MT_sim_Seq_wall.mlx* in (https://doi.org/10.5281/zenodo.8105932). The geometries of the microstructure were generated in .dxf format. For all data reported in this study, 10-20 independent simulations were executed.

## Acknowledgements

The authors would like to acknowledge N. Wingreen, D. Cohen, B. Gouveia, and P. de Souza for their valuable input during the preparation of this manuscript. Additionally, we extend our gratitude to all the members of the Petry Lab and Stone Lab for their insightful discussions throughout this project. This work is supported by the Princeton University Eric and Wendy Schmidt Transformative Technology Fund.

## Author contributions

Conceptualization: MZ, RS, HAS, SP - Methodology: MZ, RS - Investigation: MZ, RS - Visualization: MZ, RS - Funding acquisition: HAS, SP - Project administration: HAS, SP - Supervision: HAS, SP - Writing – original draft: MZ, RS, HAS, SP - Writing – review & editing: MZ, RS, HAS, SP.

## Competing interests

Authors declare that they have no competing interests.

## Supplementary Materials

**Fig. S1.**
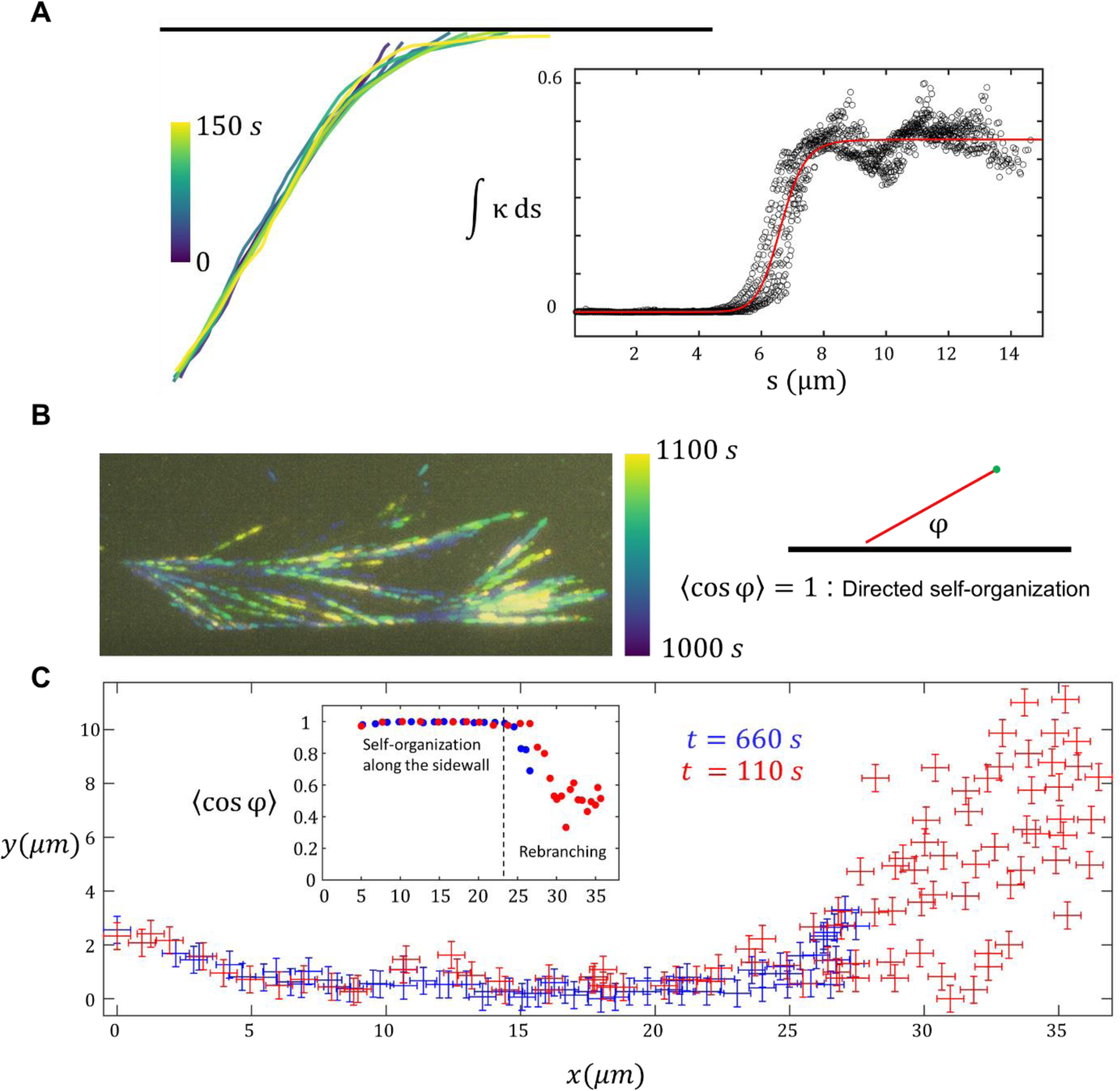
MT bending and branching along the sidewall. **(A)** MT shape when it contacts the sidewall, bends, and continues to grow along the sidewall (**left**). Relationship between MT curvature and arc length (**right**). MT bending occurs specifically at the contact point and within a span of less than 2 μm. **(B)** MT branching after self-organization along the sidewall. Sequential frames were captured at a rate of 1 fps and color-coded based on the elapsed time. **(C)** EB1 spots observed at two different time points. By probing the 〈cos(ϕ)〉 parameter, we quantified the directed self-organization of MTs along the sidewall (inset). The 〈cos(ϕ)〉 value decreases with branching, indicating that a single sidewall cannot sustain directed self-organization over long distances and periods of time.

**Fig. S2.**
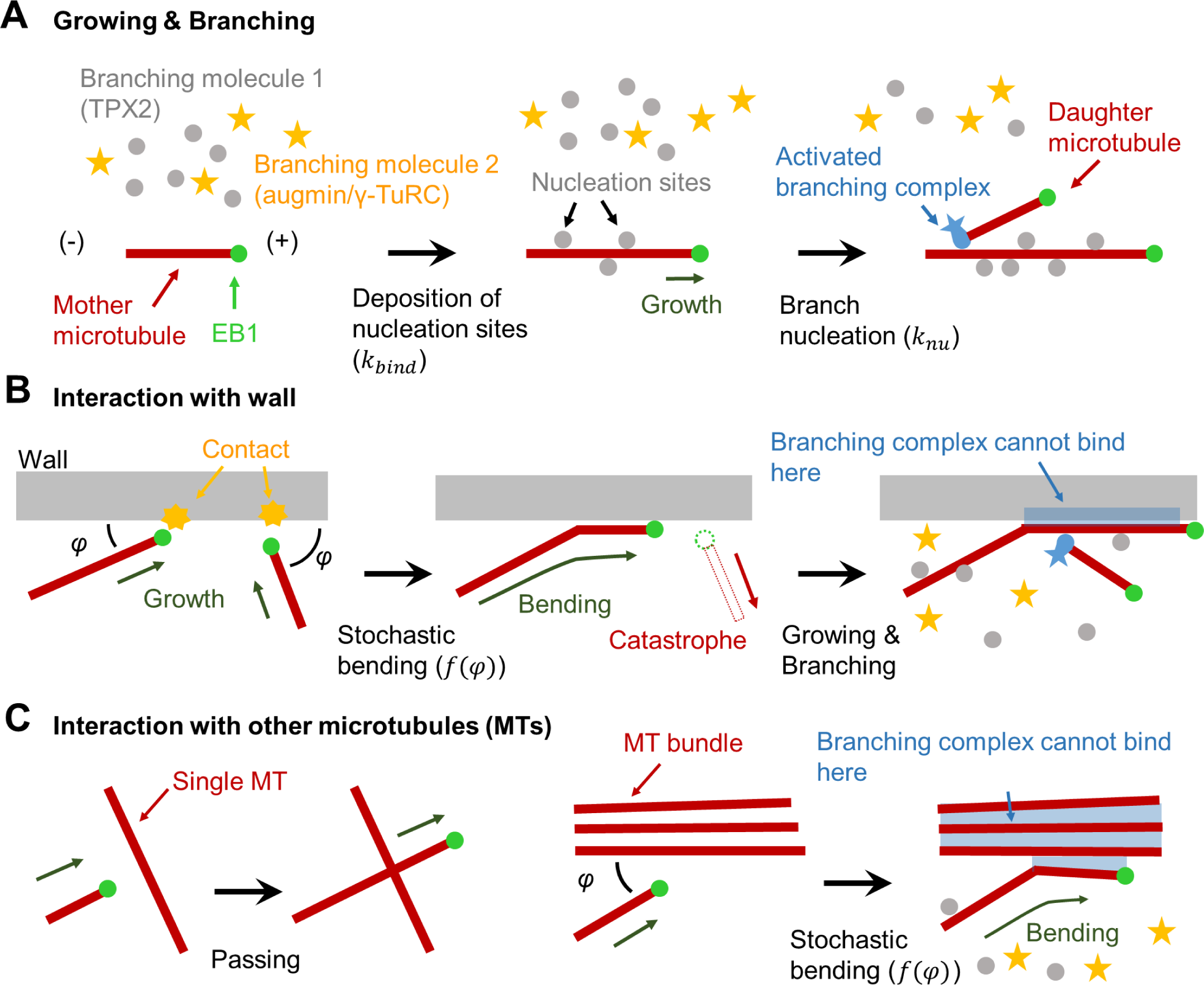
Stochastic model for self-organization of branching microtubule networks. **(A)** Schematic representation of growing and branching of MTs, benchmarking the sequential reaction model (Thawani *et al.* 2019). MT (red line) with a fixed minus end grows at a constant rate toward the plus end, denoted by EB1 (green circle). Free, inactive branching molecule 1 (gray circles) binds to an existing MT to create nucleation sites. Subsequent binding of branching molecule 2 (yellow stars) results in nucleation of a new daughter MT upon this branching complex (blue marks). **(B)** MT growth dynamics near the wall depends on their incidence angle (ϕ) with respect to the wall. Based on the probability of directed growth (*f*(ϕ)) in Fig. 2C, MTs exhibit stochastic bending towards the direction of the wall, while unbent MTs undergo catastrophe. It is hypothesized that the space for the branching complex to bind is limited between the directed MT and the wall, resulting in nucleation of a new MT only in the direction opposite to the wall. **(C)** MT growth dynamics near other MTs depends on the concentration of MTs near its plus end. When a growing MT encounters another single MT, it continues to grow without hindrance. However, a MT behaves like it encounters a wall when it encounters an MT bundle. The nucleation of new MTs is restricted within MT bundles due to the existing MTs occupying the space required for the binding of branching complexes.

**Fig. S3.**
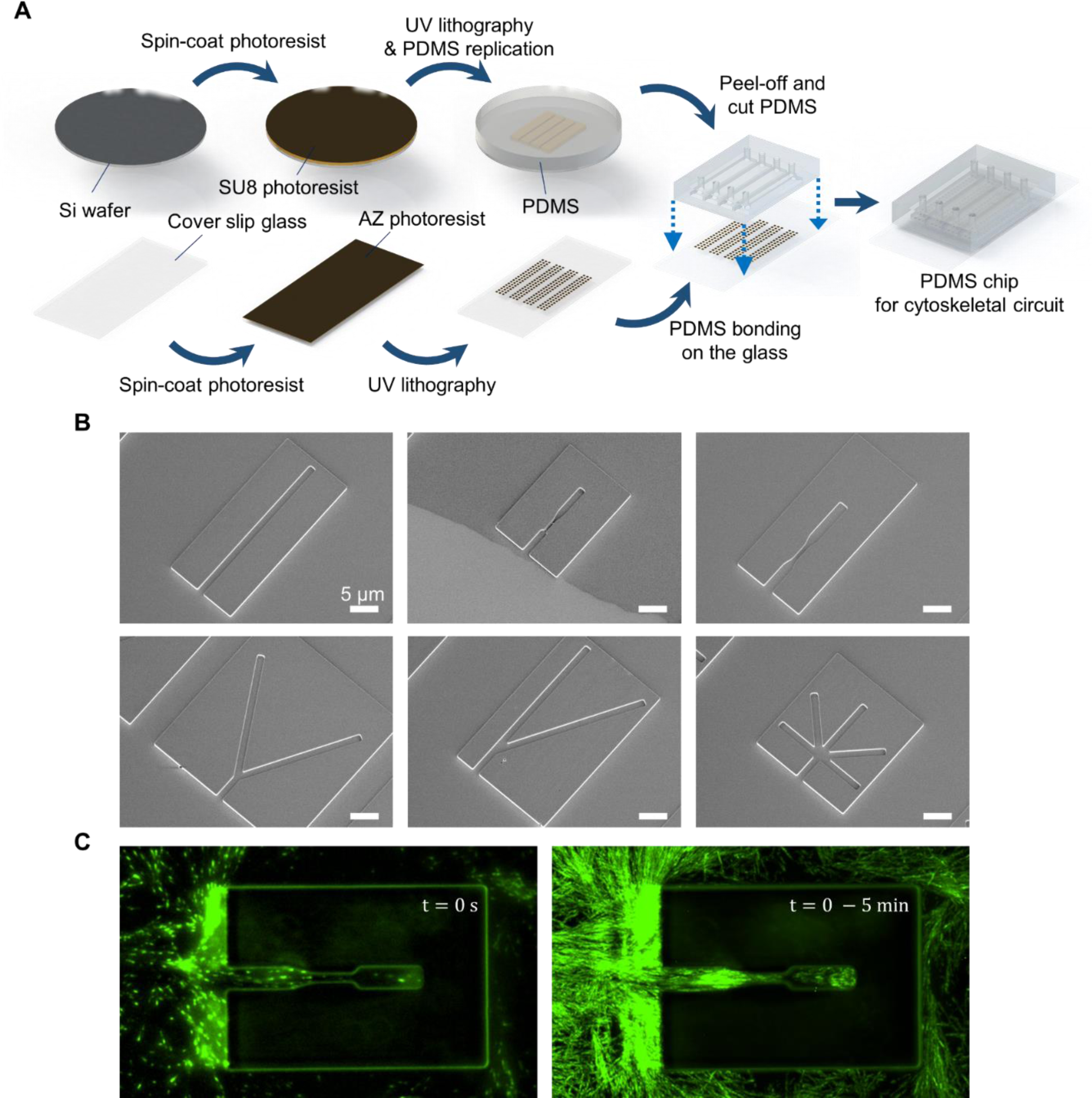
Fabrication of cytoskeletal circuit. **(A)** Schematic of the fabrication process. PDMS microfluidic chamber with microstructures on the bottom surface of the chamber. **(B)** Scanning electron microscopy images of the microstructures. **(C)** Self-organization of branched MT networks and their entry to the microstructure from side openings. **Left:** EB1 spots and **Right:** EB1 spots overlayed over 5 mins. The height of the microstructure was chosen to be 4 μm to isolate MTs inside the microstructure and allow their entry only from the side opening.

**Fig. S4.**
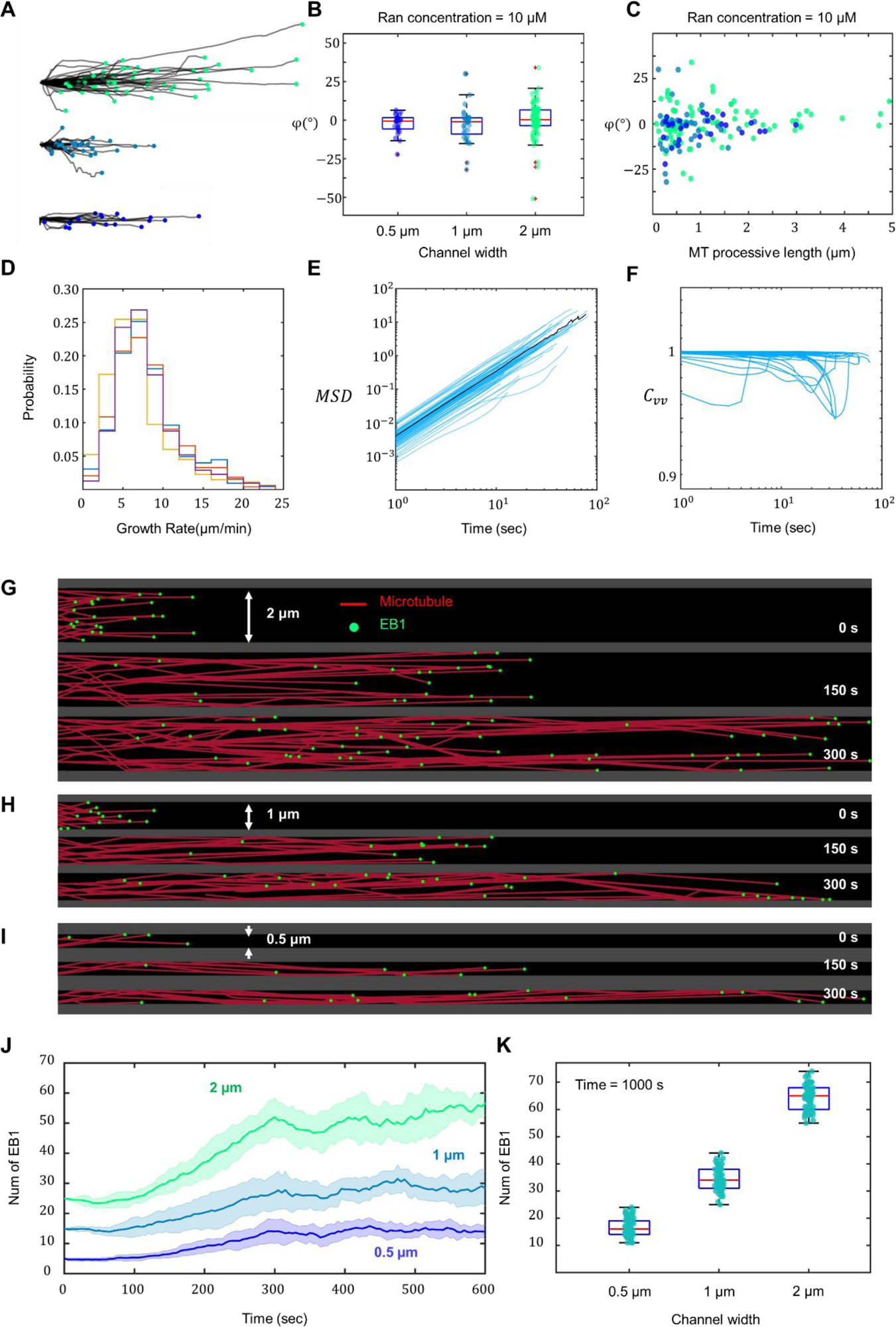
Experimental characterization and simulation of growing MTs in straight channels. **(A)** MT growth trajectories observed in channels with widths of 2 μm, 1 μm, and 0.5 μm. **(B)** MT growth angles measured in channels with widths of 2 μm, 1 μm, and 0.5 μm. **(C)** Relationship between MT growth angle and processive length (length of growth trajectories). The angle is defined relative to the channel’s long axis. **(D)** Histogram illustrating the MT growth rate at four different time points. **(E)** Mean squared displacement (MSD) analysis of growing MTs. The power-law coefficient of the MSD is approximately 1.9, indicating ballistic growth along straight lines. **(F)** Velocity correlation function of growing MTs, demonstrating the negligible loss of directionality during the growth process. **(G** to **I)** Branching growing MTs self-organize inside the channel with (A) 2 μm, (B) 1 μm and (C) 0.5 μm widths. **(J)** The number of simulated EB1 spots (growing MTs) increases over time and reaches a plateau determined by the channel width. **(K)** Number of simulated MTs in the channel according to channel width. To match the simulation results of (J) with the experimental results (Fig. 2E), 25, 15, and 5 MTs with randomized growth directions were initiated to grow from the left to the right in channels of 2 μm, 1 μm, and 0.5 μm widths, respectively, and no new MTs were added thereafter.

**Fig. S5.**
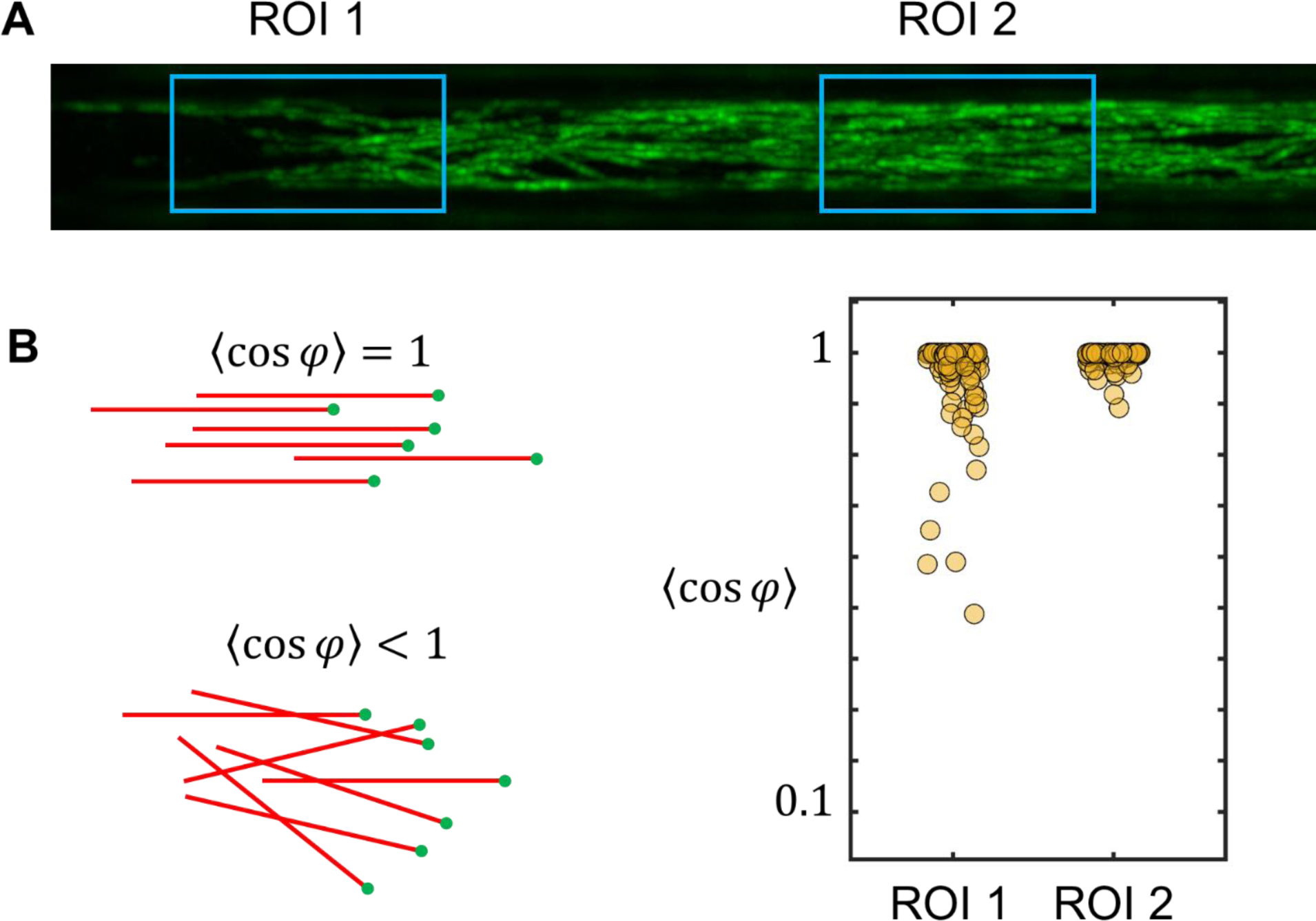
MT self-organization in a straight channel. **(A)** Overlay of frames showing EB1 spots captured at a rate of 1 fps. By probing the 〈cos(ϕ)〉 parameter, we calculated the average growth direction within two distinct regions of interest (ROIs). **(B)** As MTs self-organize within the straight channels, they exhibit a higher degree of alignment relative to the channel’s sidewalls.

**Fig. S6.**
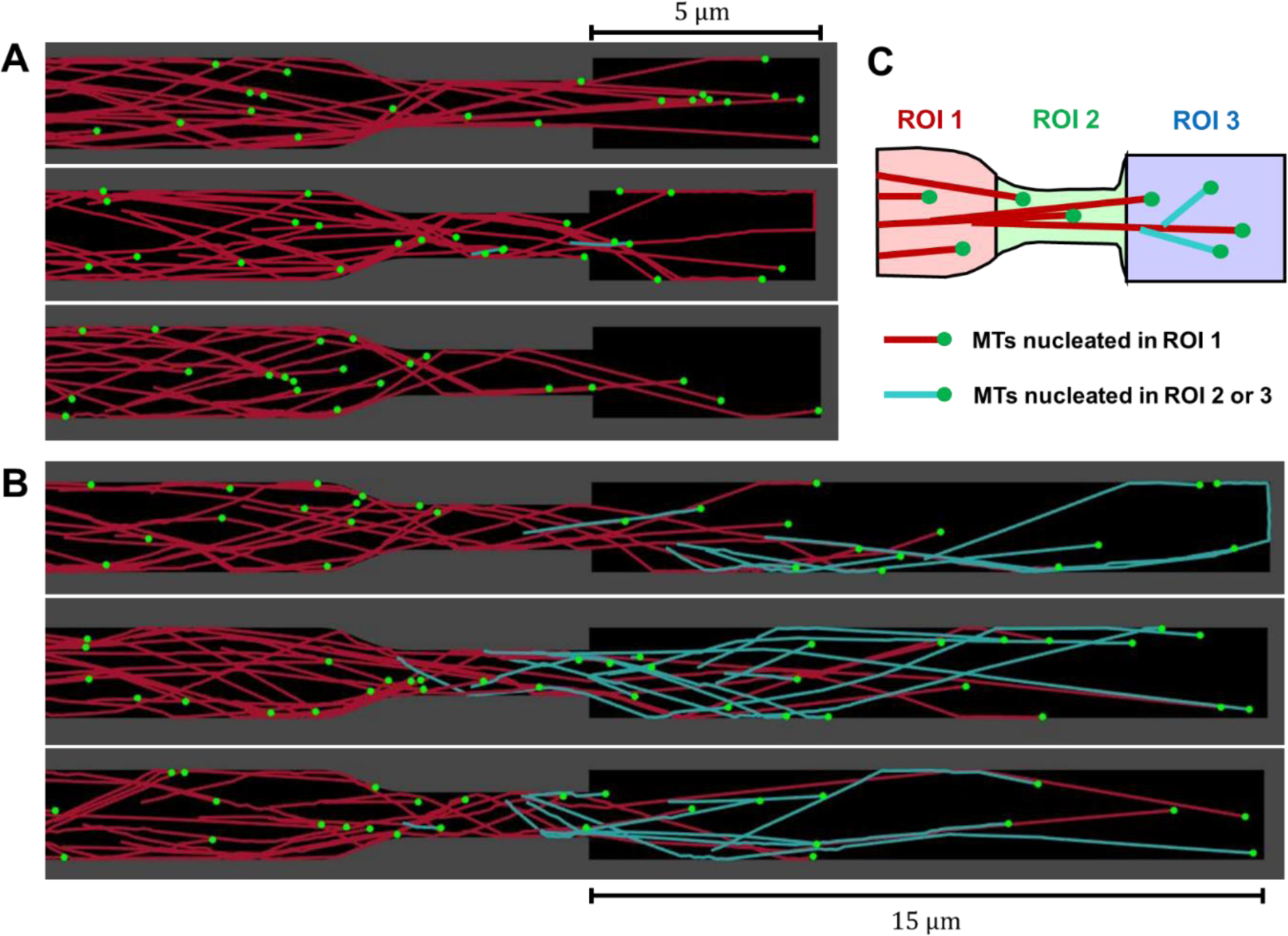
Simulation of adaptive self-organization of MTs through channels with varying width. **(A)** Schematic of self-organizing MTs inside the channel with varying width. The red-colored MTs represent the MTs nucleated in ROI 1, while the cyan-colored MTs represent the MTs nucleated in ROI 2 or 3. **(B)** In the three independent simulation results with a length of 5 µm for the third component, the cyan-colored MTs are rarely observed. **(C)** In three independent simulations with the third component length of 15 µm, the nucleation of multiple cyan-colored MTs was observed. All simulations in (B) and (C) had the same initial number of MTs with randomized growth directions (*N* = 25).

**Fig. S7.**
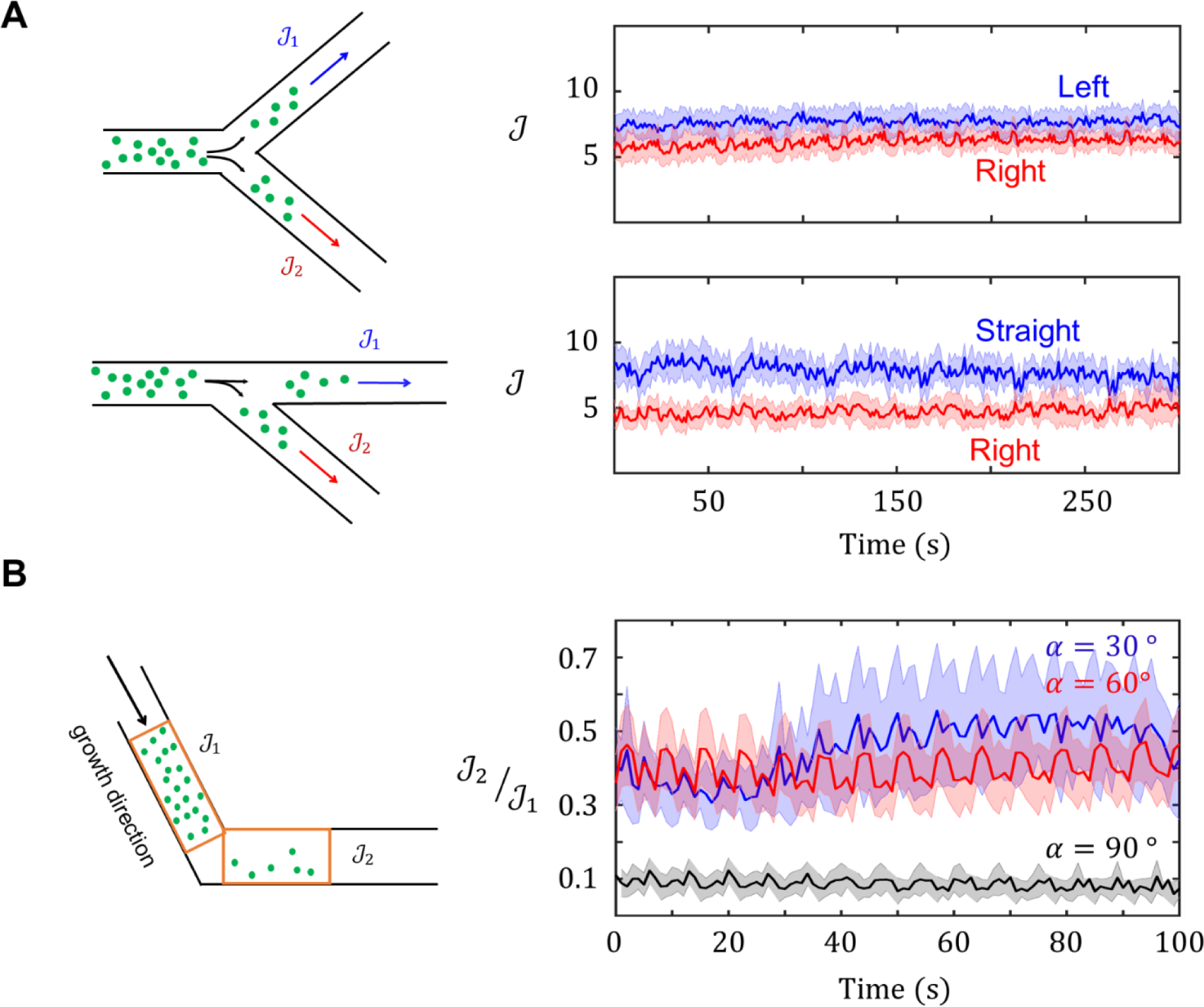
Experimental characterization of microtubule (MT) flux during divisions and turns. **(A)** MT flux observed in symmetric and asymmetric bifurcations. **(B)** MT flux measured before and after turns at different angles.

**Fig. S8.**
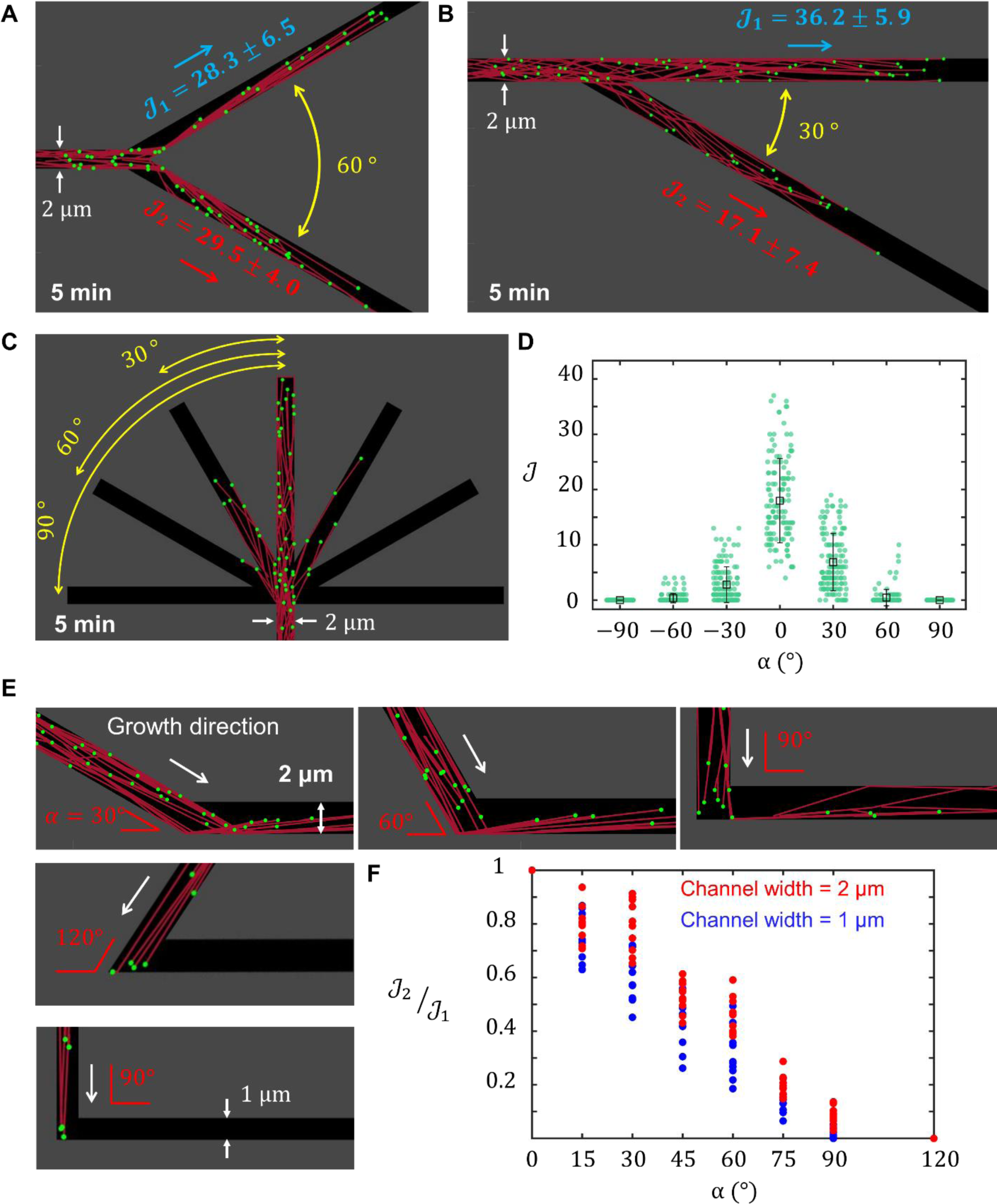
Simulation of self-organization of branched MTs network through geometrical divisions and turns. **(A** and **B)** Division of branching MTs network through (A) symmetric and (B) asymmetric bifurcations. The measured average number of EB1 spots per second were denoted as ℐ_1_ and ℐ_2_ to quantify the division process in the simulation. **(C)** Simulated MT network divisions through a hepta-furcated geometry. **(D)** The average number of EB1 spots per second in each side channel of hepta-furcated geometry. All simulations in (A) to (C) had the same initial number of MTs with randomized growth directions (*N* = 25). While the numbers of MTs in the simulations did not precisely correspond to those in the experiments, the division of branching MT networks in the simulations showed a similar trend to that observed in the experimental results. **(E)** Bending of branching MTs network through turns with different angles in the channel. **(F)** The average number of simulated EB1 spots per second before and after the turn for various angles (0 to 120°) and sizes (1 μm and 2 μm). In all simulations, 25 and 10 MTs with randomized growth directions were initiated to grow in channels of 2 μm and 1 μm width, respectively.

**Fig. S9.**
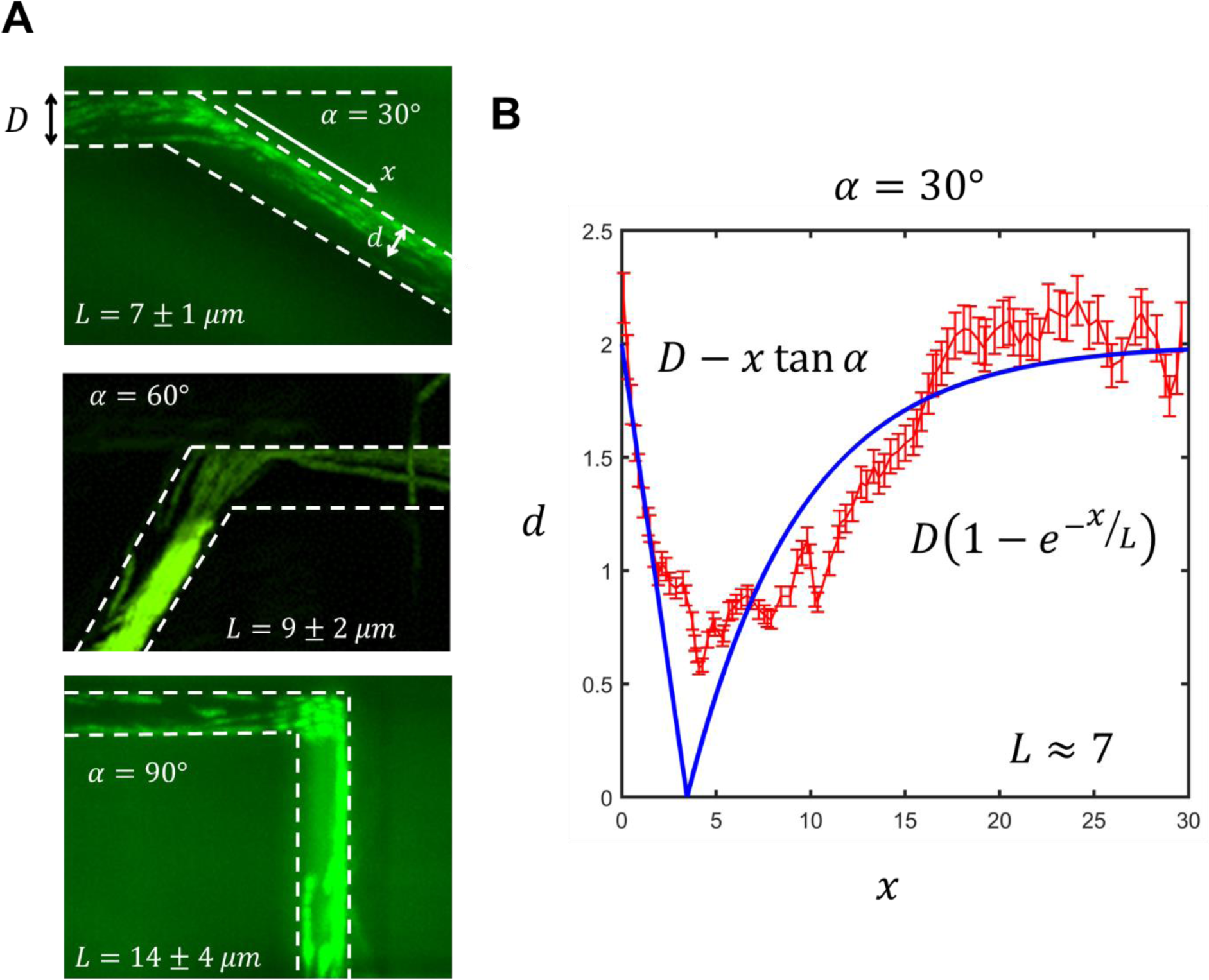
Temporary accumulation of microtubules (MTs) along a sidewall. **(A)** Overlay of frames showing EB1 spots captured at a rate of 1 fps over a 5-minute duration. **(B)** By measuring the light intensity of EB1s at various positions (indicated in red) and fitting a function (represented by the blue curve), we were able to estimate the channel length necessary for the redistribution and even filling of MTs within the channel width.

## Movie legends

**Movie S1. Directed self-organization of branching MTs along a physical boundary.** Self-organization of branching MTs without (left) and with (right) physical boundary. The scale bar is 5 μm.

**Movie S2. Branched MT networks approaching the side wall with a high incidence angle.** MTs approaching the sidewall perpendicularly shrink upon contact and do not bend. The inset in the top right corner is a 2× magnified view of the original image.

**Movie S3. Branching of MTs along the sidewall.** Branching of new MTs after directed self-organization along the sidewall: (Left) experiment and (Right) simulation.

**Movie S4. Self-organization of branching MTs within straight channels.** Self-organization of branched MT networks within straight channels with 0.5 μm (top) and 1 μm (bottom) widths: (Left) experiment and (Right) simulation. The scale bar is 1 μm.

**Movie S5. Self-organization of branched MT networks within channels featuring varying width.** Adaptive self-organization of MTs within channels with varying width: (Left) experiment and (Right) simulation. The channel width varies between 1 and 2 μm, and the left end of the channel is blocked by a wall. ROI1 to 3 are defined as in Fig. 3A and Fig. S7A.

**Movie S6. Division of branched MT networks through bifurcated and hepta-furcated channels.** Division process of branching MTs through asymmetric and symmetric bifurcation, and hepta-furcated channel with a width of 2 μm: (Top) experiment and (Bottom) simulation.

**Movie S7. Self-organization of branched MT networks within channels featuring turns.** Branching MTs within 2 μm-width channels featuring 30°, 60°, and 90° turns: (Top) experiment and (Bottom) simulation.

**Movie S8. Biased division of branching MTs.** A turn before bifurcated channel biases the division of branching MTs: (Left) experiment and (Right) simulation. A left turn before symmetric division guides MTs growth toward the right arm of the bifurcation.

**Movie S9. MT diode.** MT diodes regulate the growth direction of MTs: (Top) experiment and (Bottom) simulation. MTs self-organize from both ends of the channel towards the opposite side. The geometry of the MT diode enables forward growth while hindering growth in the reverse direction.

## Notes

### Competing Interest Statement

The authors have declared no competing interest.

